# Mapping and phasing of structural variation in patient genomes using nanopore sequencing

**DOI:** 10.1101/129379

**Authors:** Mircea Cretu Stancu, Markus J. van Roosmalen, Ivo Renkens, Marleen Nieboer, Sjors Middelkamp, Joep de Ligt, Giulia Pregno, Daniela Giachino, Giorgia Mandrile, Jose Espejo Valle-Inclan, Jerome Korzelius, Ewart de Bruijn, Edwin Cuppen, Michael E. Talkowski, Tobias Marschall, Jeroen de Ridder, Wigard P. Kloosterman

## Abstract

Structural genomic variants form a common type of genetic alteration underlying human genetic disease and phenotypic variation. Despite major improvements in genome sequencing technology and data analysis, the detection of structural variants still poses challenges, particularly when variants are of high complexity. Emerging long-read single-molecule sequencing technologies provide new opportunities for detection of structural variants. Here, we demonstrate sequencing of the genomes of two patients with congenital abnormalities using the ONT MinION at 11x and 16x mean coverage, respectively. We developed a bioinformatic pipeline - NanoSV - to efficiently map genomic structural variants (SVs) from the long-read data. We demonstrate that the nanopore data are superior to corresponding short-read data with regard to detection of *de novo* rearrangements originating from complex chromothripsis events in the patients. Additionally, genome-wide surveillance of SVs, revealed 3,253 (33%) novel variants that were missed in short-read data of the same sample, the majority of which are duplications < 200bp in size. Long sequencing reads enabled efficient phasing of genetic variations, allowing the construction of genome-wide maps of phased SVs and SNVs. We employed read-based phasing to show that all *de novo* chromothripsis breakpoints occurred on paternal chromosomes and we resolved the long-range structure of the chromothripsis. This work demonstrates the value of long-read sequencing for screening whole genomes of patients for complex structural variants.

## Introduction

Second-generation DNA sequencing has become an essential technology for research and diagnosis of human genetic disease. Sequencing of human exomes has resulted in the identification of genes involved in Mendelian disorders ^1^, while whole-genome sequencing has revealed that a myriad of diseases is caused by genetic changes that can occur both within genes as well as in the noncoding genome ^2^. As a result, genome sequencing has seen rapid adoption in clinical decision making as a complete picture of a patient’s unique mutation profile enables personalization of treatment strategies ^3,4^.

Robust methods to detect structural variants (SVs) in human genomes are essential, as SVs represent an important class of genetic variation that accounts for a far greater number of variable bases than single nucleotide variations (SNVs) ^5^. Moreover, SVs have been implicated in a wide range of genetic disorders ^6^.

A particularly revolutionary development in genome sequencing is the usage of protein nanopores to measure DNA sequence directly and in real time ^1,7^. The first successful implementation of this principle in a consumer device was achieved in 2014 by Oxford Nanopore Technologies with the introduction of the MinION ^8^. The MinION can sequence stretches of DNA of up to hundreds of kilobases in length, which already resulted in the sequencing of the genomes of several organisms^9,10^.

An important and natural application of the long reads produced by nanopore sequencing is identifying structural variations. Long-read sequencing is breaking ground for the discovery of SVs at an unprecedented scale and depth ^11^. The first success has been achieved using the Pacific BioSciences SMRT long-read sequencing platform ^12,13^ and alternative methods including BioNano ^14^ and 10X Genomics ^15^. While short-read next-generation sequencing data rely on multiple (often) indirect sources of information in order to accurately identify SVs, structural changes can be directly reflected in long read data.

In this work, we demonstrate, for the first time, the sequencing of the whole diploid human genome on the MinION sequencer. We sequenced the genomes of two patients with congenital disease resulting from chromothripsis at 11-16X coverage depth. We employ a novel computational pipeline to demonstrate the possibility of using nanopore reads to detect *de novo* complex SV breakpoints at high sensitivity. The long reads from the MinION allowed efficient phasing of genetic variations genome wide (SNVs as well as SVs) and enabled us to resolve the long-range structure of the chromothripsis in the patients. Moreover, we identify a significant proportion of SVs that are not detected in short-read Illumina sequencing data of the same patient genomes.

These results highlight the feasibility to sequence clinical human samples in real-time on a low-cost device. Because nanopore-based sequencing requires almost no capital investment and current devices have a very small footprint, mainstream adoption of these sequencers has the potential to fundamentally change the current paradigm of sequencing in centralized centers.

## Results

### Long-read whole genome sequencing of patients genomes using the MinION

As a first step toward real-time clinical genome sequencing, we evaluated the use of the MinION device to sequence the genomes of two patients with multiple congenital abnormalities ^16^, denoted hereforth as Patient1 and Patient2 respectively.

We extracted DNA from patient cells and sequenced this on the MinION. For Patient1, we used R7, R9 and R9.4 pore chemistries (**Supplementary Table 1**) generating a total of 8.2M template sequencing reads from 122 sequencing runs. For Patient2, we exclusively used R9.4 runs and performed only 13 runs (1.89M reads), which required approximately five days of sequencing on seven parallel MinION instruments at a cost of around $7,000. This demonstrates the feasibility of sequencing whole human genomes at >10X coverage with the latest R9.4 chemistry (**Supplementary Figure 1**). 82.1% (Patient1) and 98.9% (Patient2) of these reads could be mapped to the human reference genomes and were useful for further analyses. Read lengths were highly variable for Patient1, as a result of differences in library prep methods, with a mean of 6.9kb for template reads, while for Patient2 we reached an average of 16.2kb with consistent read length distributions for each of the 13 runs (**Supplementary Figure 2**).

Raw sequencing data were transformed into FASTQ format using Poretools and sequence reads were mapped to the human reference genome (GRCh37) using LAST ^17^. We uniquely aligned 99% of R7/R9 2D reads or R9.4 1D reads flagged as ‘passed’ after EPI2ME basecalling, while this dropped to 55% for R9-based ‘failed’ 2D reads (**Supplementary Figure 3**). We evaluated the mapping accuracy by calculating the percentages of identical bases between mapped reads and the reference genome (PID). We observed a mean PID of 90% for R7 2D and R9 2D, 85% for R9 template and 89% for R9.4 template reads based on LAST mapping (**Supplementary Figure 4**).

We obtained a mean coverage depth of 16X and 11X for Patient1 and Patient2, respectively (**Supplementary Figure 5**). Coverage was lower in regions with extreme GC content, yet this effect was significantly much less pronounced than for Illumina sequencing of the same genomes (**Supplementary Figure 6**) ^12^. This finding was confirmed by analysis of k-mer distributions of nanopore and Illumina data (**Supplementary Figure 7**). We noted that while the nanopore reads marked as ‘fail’ show overall systematic sequencing biases regarding coverage distribution, the quality of the aligned fraction is comparable to the ‘pass’ reads. We therefore included the ‘fail’ data of Patient1 that was successfully retrieved through alignment, in all subsequent analysis.

### Resolving de *novo* genomic rearrangements using long-read data

Both patients have complex phenotypes involving dysmorphic features and mental retardation, likely caused by their *de novo* complex chromosomal rearrangements, which were karyotypically defined as 46,XX,ins(2;9)(q24.3;p22.1p24.3)dn (Patient1) and 46,XY,t(1;9;5)(complex)dn (Patient2) ^16^.

We evaluated the performance to detect the breakpoints underlying the complex *de novo* karyotypes of Patient1 and Patient2 using nanopore sequencing data, at this medium coverage. Both patients have already been described in recent work, in which Illumina sequencing was used to map the rearrangement breakpoints, which is the current gold-standard method for routine genome-wide SV mapping in patient genomes ^16,18^. For Patient1, we augmented the previously described data by performing Illumina HiSeq X data for both parents. We performed SV calling with Delly ^19^ and Manta ^20^ on the Illumina data from Patient1 and its parents. By integrating SV calls from Delly and Manta and removing calls that were also identified in one or both parents, we obtained a set of 44 putative *de novo* SV breakpoints, 40 of which formed a complex genomic rearrangement, as described previously^16^. These 40 breakpoints could be verified by orthogonal breakpoint assays using PCR and MiSeq sequencing (**Supplementary Table 2**). The breakpoints cluster within regions of chromosomes 2, 7, 8 and 9 and are the result of a complex shattering and reassembly process, known as chromothripsis ^21,22^ (**Figure 1A**).

**Figure 1.**
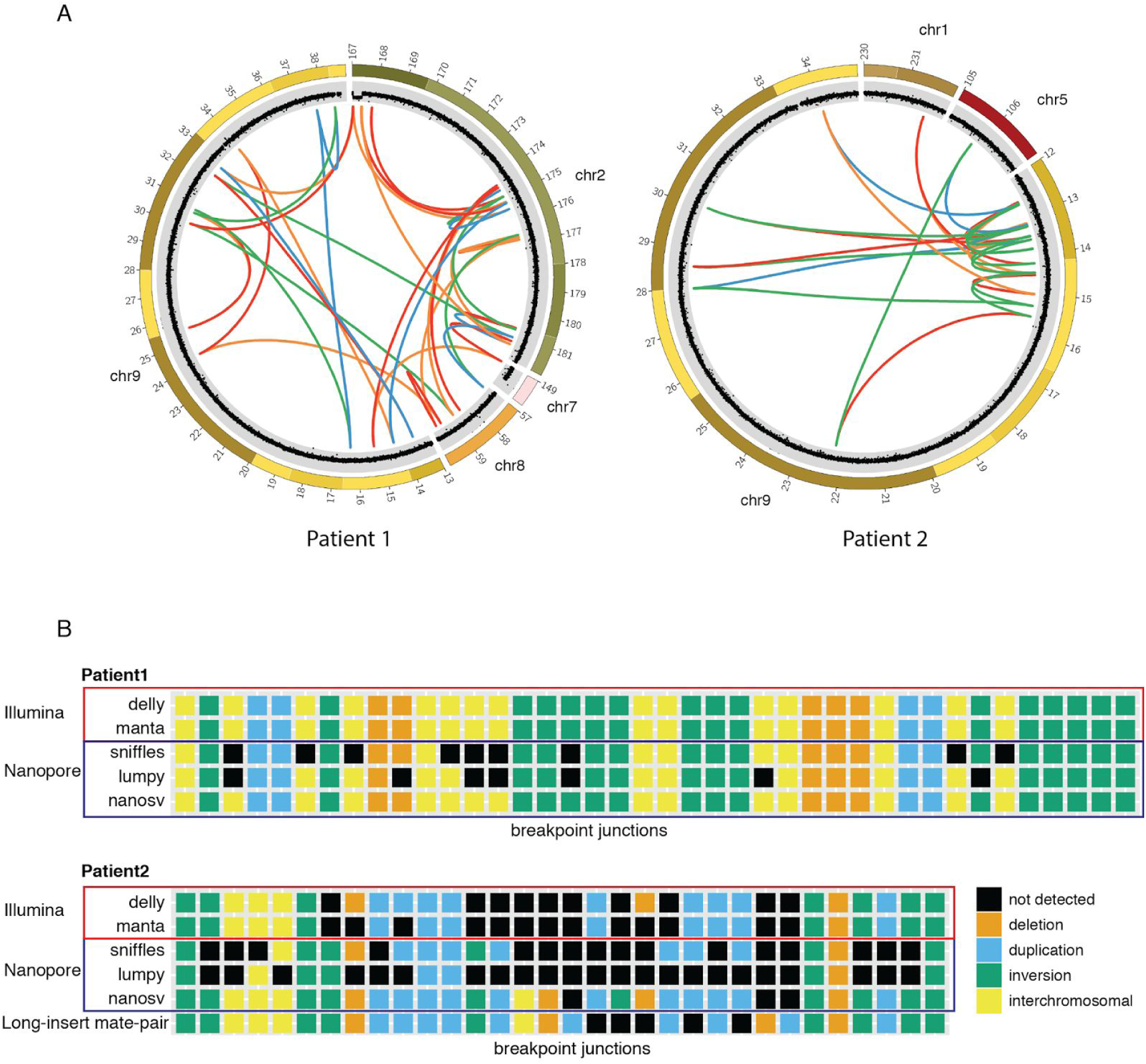
De novo breakpoint junctions involved in complex chromosomal rearrangements of Patient1 and Patient2.

(A) Circos plots for Patient1 and Patient2 respectively. For Patient1, we took the set of 40 validated *de novo* breakpoint junctions obtained by Illumina whole-genome sequencing of the patient and its parents. For Patient2, we depicted the breakpoint-junctions as published recently ^52^. The outer ring of the circos plot shows the chromosome ideogram and the inner ring shows the copy number changes as revealed by FREEC ^53^ analysis of Illumina whole genome sequencing data for both patients. Colored lines indicated breakpoint-junctions. Blue: tail-to-head, green: head-to-tail, red: head-to-head, yellow:tail-to-tail. (B) SV genotyping comparison across the chromothriptic breakpoint junctions, between Illumina Hiseq data and Nanopore data, using various tools tested. The x-axis represents different breakpoint-junctions and the y-axis shows different SV calling methods and datasets. The individual breakpoint-junctions are indicated by colors specifying the type of breakpoint junction.

For Patient2, there were 29 SVs underlying the complex *de novo* karyotype as based on the previously described breakpoint-junctions, which were detected using long-insert mate-pair sequencing and revealed a complex chromothripsis rearrangement involving chromosomes 1, 5 and 9 (**Figure 1A**, **Supplementary Table 2**) ^16^.

To enable SV detection in nanopore long-read sequencing data, we developed a new bioinformatic tool, NanoSV, tailored to nanopore data. NanoSV uses split-read mapping (obtained from LAST alignment) as a basis for SV discovery (**Methods, Supplementary Figure 8**), and supports discovery of all defined types of SVs (**Supplementary Figure 9**). We used NanoSV to detect SVs in the nanopore sequencing data generated for Patient1 and Patient2. To benchmark NanoSV against other SV callers, we tested Lumpy ^23^ and Sniffles ^24^ for the Nanopore data and Manta and Delly for the Illumina data. All other callers (i.e. except NanoSV) require BWA alignments as input.

For Patient1, we reached 100% sensitivity with regard to detection of the 40 validated breakpoint-junctions. Conversely, we identified 33 (83%) and 31 (78%) of the 40 *de novo* breakpoint junction in the call sets from Lumpy and Sniffles, respectively (**Figure 1B**). For Patient2, NanoSV detected 24 of the 29 previously described breakpoint-junctions. We investigated further why five variants were missed, using Sanger sequencing of PCR products of the respective breakpoint-junctions. We found that two out of the five previously published breakpoint-junctions represent a complex combination of more than two joined segments (**Supplementary Figure 10**, **Supplementary Table 2**). These short segments were likely missed by the long-insert jumping libraries that were used in the previous work ^16^. Based on validation by Sanger sequencing, we retrieved total of 32 chromothripsis breakpoint-junctions in Patient2 and 29 (91%) of these were detected using NanoSV (**Figure 1B**). The three remaining junctions were missed by nanopore sequencing because of the low sequencing coverage, as we could observe split read mappings supporting each of these junctions. Sniffles and Lumpy, detected 16 (50%) and 9 (28%) of the 32 breakpoints-junctions in the nanopore data from Patient2, respectively; Manta and Delly detected 19 (59%) and 22 (69%) of the 32 breakpoint-junctions respectively, in the short-insert Illumina data of Patient2.

### Unraveling the long-range structure of the chromothriptic region

In earlier work it has been suggested that germline chromothripsis originates on paternal chromosomes ^21^, but this has been inferred from only a few breakpoint-junction sequences or deleted segments. A thorough validation of the conjecture that the origin of chromothripsis is exclusively paternal is lacking. Furthermore, the structure of the chromothripsis rearrangements has been inferred from the patterns of breakpoint-junctions, under the assumption that the chromothripsis breakpoint-junctions occur on a single haplotype ^21,22,25^.

We developed a bioinformatic pipeline to augment genome-wide genetic SNP phasing with nanopore read-based phasing of SVs (**Methods**). In a first step we obtained 1.7M heterozygous SNPs from Patient1, that were called from Illumina sequencing data and trio-phased using GATK PBT ^26^ and Patient1’s parents’ genotypes (alternatively, statistical phasing from publicly available reference panels may be used). Subsequently, each nanopore read was assigned phase based on a phasing score that takes into account the content and number of overlapping phase-informative SNPs (**Methods**). Per chromothripsis breakpoint-junction, we obtained between 2 and 11 break-supporting nanopore reads and 85% (195/228) of these overlapped on average of 9.8 phase-informative heterozygous SNPs. Additionally, we similarly phased the Nanopore reads that spanned but did not support the breakpoint junctions (reference reads). This analysis demonstrated that all 40 *de novo* chromothripsis breakpoint-junctions are unequivocally of paternal origin (**Figure 2**), whereas non-breakpoint-spanning reads at the same loci are of maternal origin. A small fraction of the reads points to an origin of some breakpoint-junctions on maternal chromosomes. These are all reads with three or less overlapping phase-informative SNVs, and therefore likely represent artifacts. These results support earlier hypotheses of a paternal origin of germline chromothripsis, pointing to a breakage and repair process specific for male chromosomes occurring either during spermatogenesis or early zygotic cell divisions ^27^. We were further able to reconstruct Patient1’s affected, derivative chromosomes by following the chain(s) of breakpoint-junctions by order and orientation (**Figure 3A-B**). Such a strategy leads to a configuration of four derivative chromosomes for Patient1, each containing one centromere and two telomeric chromosome ends. The such obtained chromosomal structure matched the observed karyotype (**Supplementary Figure 11**).

**Figure 2.**
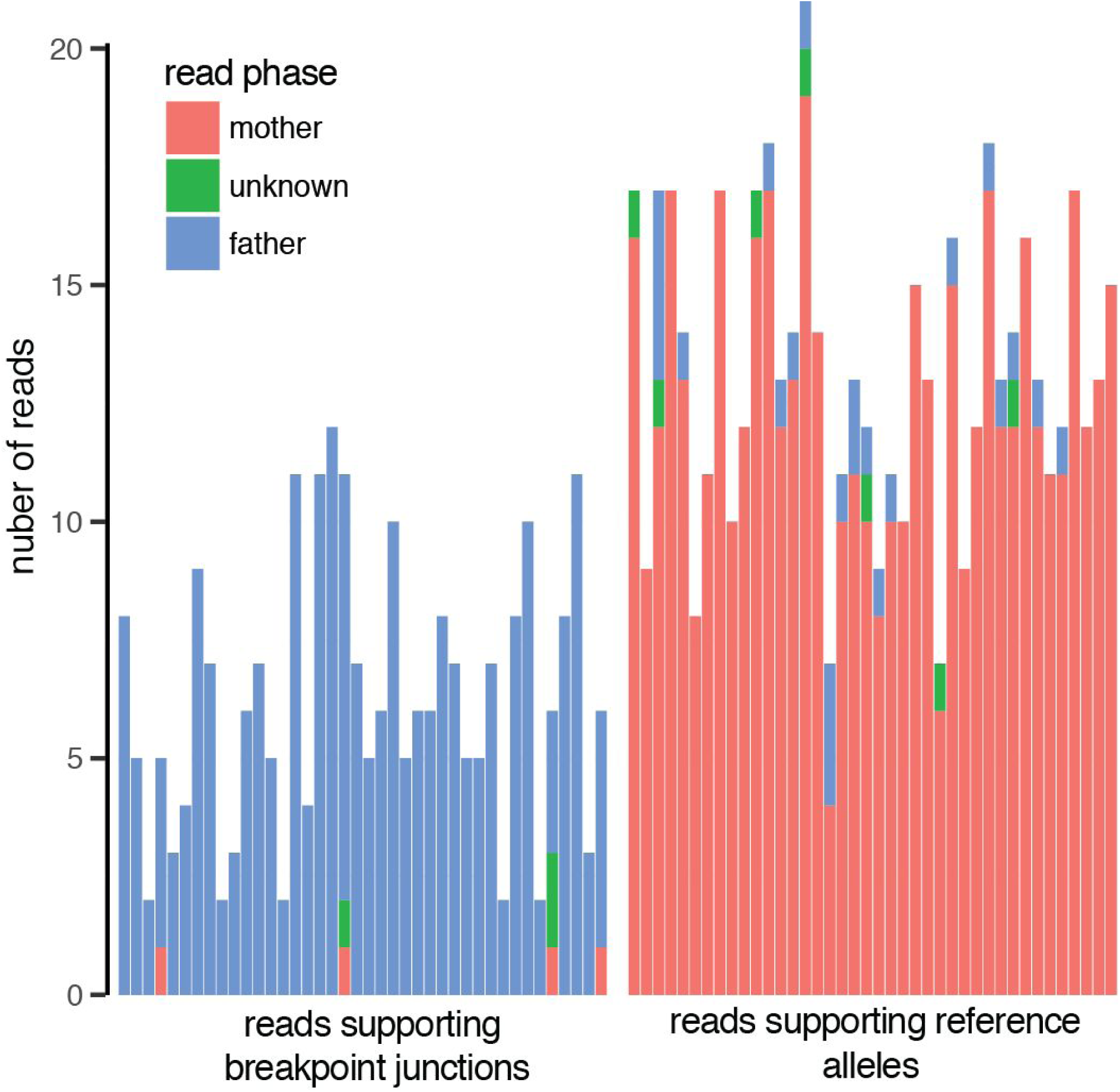
Phasing of chromothripsis breakpoint-junctions.

Bardiagram displaying phasing of nanopore reads overlapping 40 chromothripsis breakpoint-junctions in Patient1. The x-axis displays each of 40 chromothripsis breakpoint-junctions identified in Patient1, stratified by allele (alternative and reference). On the left side only reads supporting the alternative allele are depicted and on the right side reads supporting the reference allele are shown. The y-axis indicates the number of reads supporting each allele, for each of the 40 breakpoint-junctions. Legend colors indicate whether the assigned read phase was paternal, maternal or unknown.

**Figure 3.**
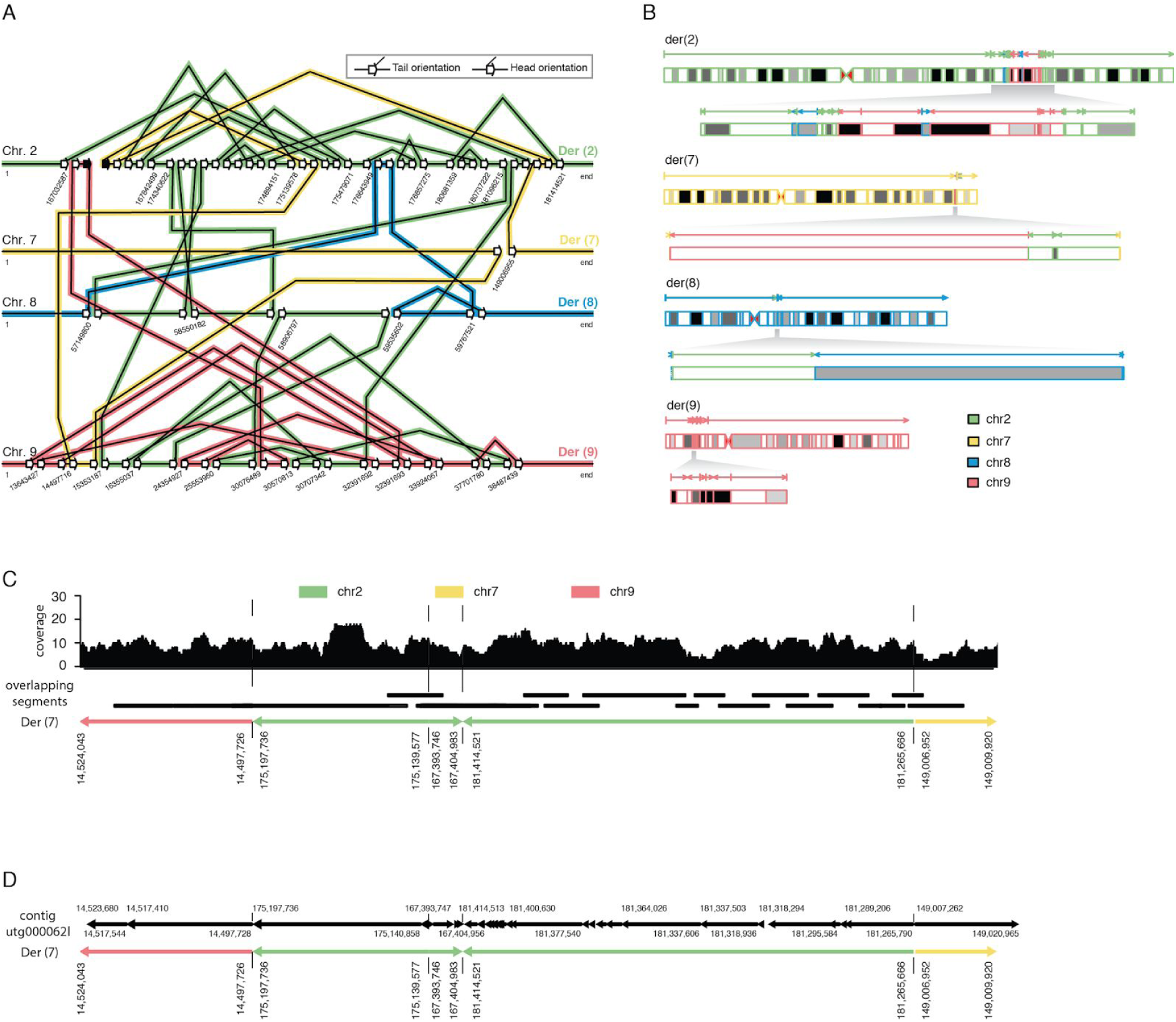
Unraveling long-range chromothripsis structure from the read data.

(A) Schematic diagram showing the patterns of breakpoint-junctions in Patient1. The human reference genomic regions that are involved in the chromothriptic event are depicted horizontally for each affected chromosome. The slanted lines connecting various reference segments represent breakpoint-junctions. The orientations of breakpoint-junctions are indicated by arrows as shown in the legend. Black (instead of open) arrows indicate the boundaries of a chromosomal deletion resulting from the chromothripsis, whereas open arrows indicate double-stranded DNA breaks. (B) Structure of chromothriptic derivative chromosomes in Patient1 as inferred from the orientations and order of breakpoint-junctions shown in panel A. (C) Reconstruction of a chromothriptic subregion of chromosome 7, involving 5 chromosomal segments. Overlapping aligned reads originating from Patient1’s paternal haplotype were used. Nanopore reads that are instrumental for segment connectivity are indicated by black bars. The coverage track has been generated from all paternal reads mapping to the respective chromosomal segments. The underlying derived chromosome’s structure is illustrated on the bottom. (D) Haploid assembly of the chromothriptic region of Patient1. A 469 kb contiguous assembled sequence (utg000062l) spans, through 54 segments that align back to the reference genome, the same chromothripsis subregion illustrated in panel C. The fragmentation of the assembly into many (54) aligned segments is expected given that Miniasm does not compute a consensus sequence.

We further wished to investigate the extent to which the derived chromosomal structure can be reconstructed from the nanopore sequencing data. We note that a much higher (Nanopore) sequencing depth is required in order to accurately reconstruct such large chromothriptic regions through diploid assembly. In order to evaluate the potential of Nanopore long-read data to facilitate future analyses, we pre-phased, as described above, all the reads that align within the chromothriptic region (i.e.: ∼40MB of genomic sequence across 4 chromosomes) and use only the reads that are known to originate from the paternal haplotype and those that cannot be assigned phase (i.e. where the two haplotypes are identical).

We built contigs first by evaluating the read overlaps from the reference alignment (**Methods**) and obtained contigs that connect between two to five chromothriptic segments, spanning up to 2MB of contiguous DNA sequence (**Figure 3C, Supplementary Figure 12**). Finally, we used Miniasm^28^ to evaluate whether such longer, local haplotype structure can be retrieved in a more easily scalable fashion (**Methods**). The whole 40MB region was assembled into 178 contigs that were subsequently aligned against the human reference genome. We identified three contigs of 241kb, 469kb and 1,217kb in size, each spanning 3 to 5 chromothriptic segments. Segment order and orientation in each of the three contigs supports the predicted chromothripsis structure (**Figure 3D, Supplementary Figure 13**).

### Genome-wide structural variation discovery from nanopore reads

Beyond very specific, targeted applications, long sequence reads present unique advantages for genome structural variation discovery in human genomes, as it has been recently shown from data generated on Pacific Biosciences platforms ^12,29^. Here, we assessed whether our Nanopore sequencing data could yield any novel SVs beyond those found in Illumina short-read sequencing of the same sample by using Patient1 as a test case. We used NanoSV to produce an initial call set of 33,592 SV breakpoint-junctions. Manual inspection of SV candidates showed that Nanopore sequencing and base-calling is very poor in regions containing homopolymer stretches, which typically lead to a collapse of the whole region into a spurious indel call. This is observed across samples, as well as in nanopore-based high depth of coverage resequencing of PCR products (**Supplementary Figure 14**). Additionally, we noted that SV calling is similarly hampered in tandem repeat regions (**Supplementary Figure 14**). Based on these observations and in order to obtain a good quality consensus call set, we discarded calls for which both ends of the candidate breakpoint-junction fall within genomic homopolymer regions or short tandem repeat stretches, and remain with a set of 8,667 SV breakpoint-junctions for Patient1.

The remaining call set of 8,667 SVs was intersected, for reference, with calls generated by two additional nanopore SV callers (Lumpy, Sniffles). Furthermore, we performed SV calling on the corresponding Illumina data of Patient1 using six tools (Pindel, Manta, Delly, FREEC, Mobster and GATK HaplotypeCaller) that are commonly used in human genome sequencing studies and which represent different methods to detect SVs from whole genome short-read Illumina sequencing data that collectively capture most classes of SVs ^19,20,30–32^. An SV is considered to be overlapping with the Illumina dataset if the Nanopore data SV call matches an SV call in any of the tools used on the Illumina data. We further considered as overlapping Illumina data (i.e. “detectable” through short read sequencing) any NanoSV called variant that can be matched within the 1,000 genomes SV and indel sites respectively (**Supplementary Figure 15**) ^33^.

We performed multiple rounds of orthogonal validation, on a random sample, spanning all SV classes and size ranges (**Methods**). The estimated precision of our 8,667 SV calls is 70% as based on 273 validation assays (193 true positives and 80 false positives). We further devised a post-calling filtering step, in order to produce a high confidence set of SV candidates. We manually curated an additional (random) subset of 83 SV calls and defined a training set of 191 true-positive and a set of 90 false-positive SV calls (**Methods**). These data were subsequently used to train a random forest classifier, aiming to filter out false positive calls. The features that are included in the model are extracted from the aligned sequencing data and are designed to be sequencing read-depth and read-length independent, such that the model be applicable to any Nanopore sequencing setting (**Methods**). Based on a model with an estimated 92% precision and 83% recall (**Supplementary Figure 16**), we obtain a final set of 4,778 SV calls for Patient1, out of which 3,265 variants are found (or detectable) in short-read sequencing data as well. The remaining 1,513 (32%) variants are exclusively found in the Nanopore long-read sequencing data. Evaluation of a set of 75 SVs that were tested for validation, but not used for training the random forest model, shows a precision of 84% on our final set of SVs. We applied the same computational pipeline to call and filter SVs from the Nanopore data of Patient2, as well as intersect with SV calls from illumina short read data. After applying the random forest model trained on Patient1 data, we obtain a set of 4,977 high confidence SV calls for Patient2, out of which 3,237 are also identified in Illumina data, while 1,740 SVs are Nanopore specific. Aggregating the data from Patient1 and Patient2 we obtain a set of 6,502 SVs, detectable by short read sequencing as well as long-read sequencing and a set of 3,253 SVs that are long-read data specific.

A comparison of the two sets of SV calls shows that Nanopore specific SVs are located at sites with a higher GC content (i.e.: than SVs also genotyped from illumina data) on average, which are typically hard to sequence with short-read technologies (**Supplementary Figure 17**). The most frequent class of SVs in the set of 9,755 predicted true-positive SVs are deletions (49%), yet the major fraction (84%) of these are also found in short-read data (**Figure 4A**). The largest fraction of Nanopore-specific variants is found for (tandem) duplications, where 60% (1,849) were not detected in corresponding Illumina data (**Figure 4C**). Visual inspection of several genomic sites containing such short duplications, revealed that limited evidence for some of these can also be observed from Illumina data, yet at a medium to high coverage of 40x none of 6 SV calling algorithms makes an alternative allele call. Similarly, we found that 34% of insertions in the size range of 51 to 200 bp are only identified in the Nanopore data. Overall, most Nanopore-specific SVs are found in the 10-200 size bin, which is a size range that is notoriously difficult for short-read data ^34^. Finally, we find that 45% of the large inversions (>10kb) are only identified in our Nanopore-specific set of SVs (**Figure 4D**).

**Figure 4.**
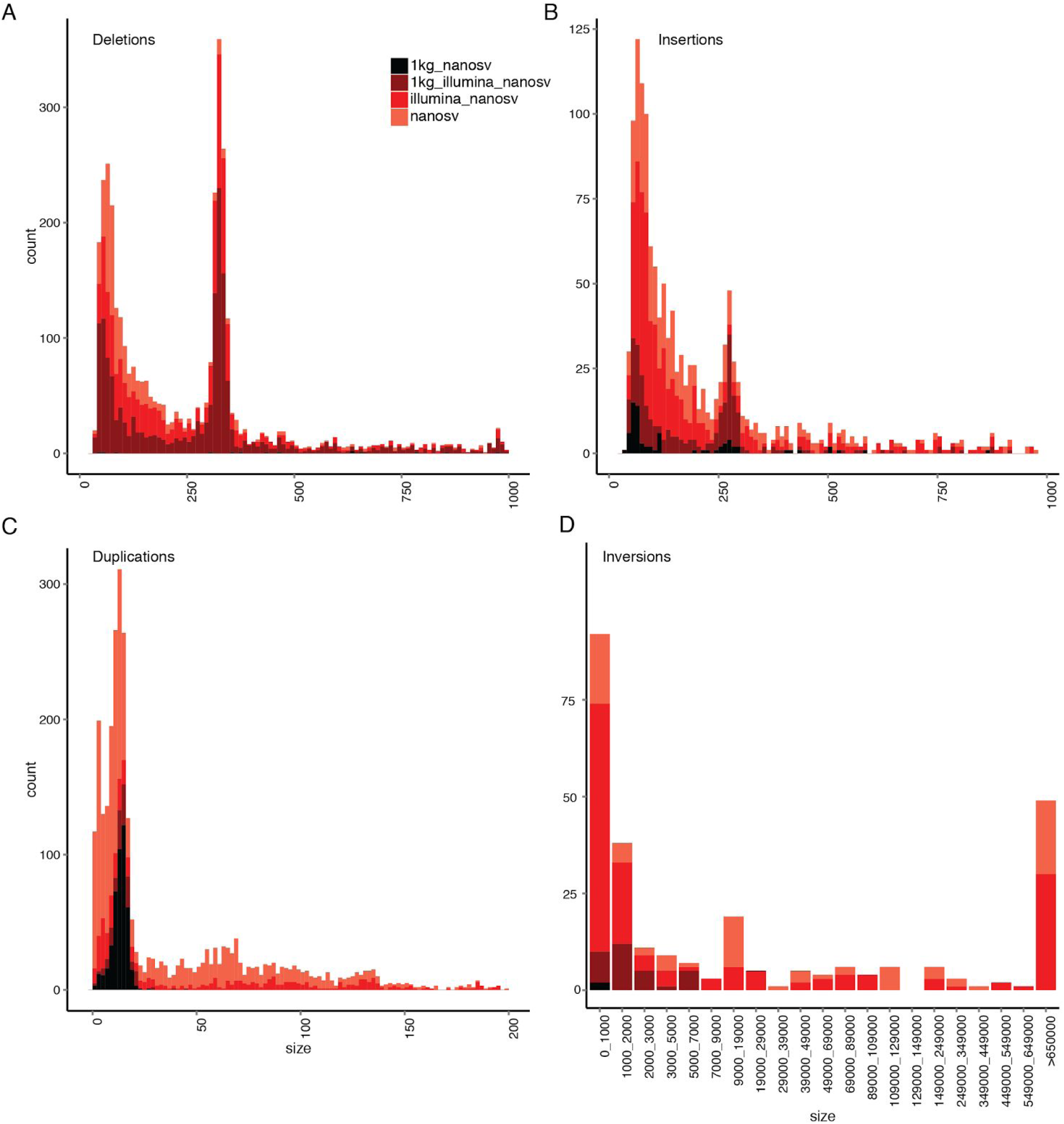
Detection of novel SVs using nanopore sequencing data.

Barplots indicating the total amount of high confidence NanoSV SV calls for Patient1 and Patient2 jointly, across different SV size bins and stratified by SV type as follows: deletions (A), insertions (B), duplications (C) and inversions (D). The NanoSV calls were intersected with SV calls from other data sources (Illumina data of Patient1 and Patient2 and 1000 Genomes phase 3 sites). For panel A and B, the x-axis was trimmed to 1000bp and a small number of variants beyond this size are not displayed in the figure. Similarly, for panel C the x-axis was limited to 200bp.

### Nanopore read-based phasing of single-nucleotide variations

Phasing genetic variation is critical for human disease studies ^35,36^. To demonstrate the potential of long-read sequencing data for direct read-based phasing of genetic variation, we employed WhatsHap, an algorithm that we recently established ^37,38^. Using WhatsHap, we phased a set of high-quality genome-wide SNVs from both patients (**Methods**) and obtained haplo-blocks with N50=126kb for Patient1 and N50=305kb for Patient2 respectively. The distribution of block lengths is shown in **Figure 5A**. We were able to establish 97.5% (96.5%) of all possible phase connections in Patient1 (Patient2), where a phase connection is defined as the relative phase between two consecutive heterozygous SNVs (**Figure 5B**). For Patient1, where Illumina sequencing data was available for the parents, we produced a ground-truth phasing by genetic haplotyping, that is, by using the SNV genotypes and the family relationship ^26^. Additionally, we phased both samples using ShapeIt2 and the 1000 Genomes phase 3 reference panel ^39^. **Figure 5C** shows pairwise comparisons of the obtained haplotypes, with switch error rates of 1.7% and 2.3% when comparing read-based and population-based phasing for Patient1 and Patient2, respectively. We observed a lower switch error rate of 1.4% between trio-based and read-based phasing, which points to a significant amount of switch errors in the population-based phasing (1.0% when comparing trio-based vs. population-based phasing). Therefore, a significant amount of differences between read-based and population-based phasing is most likely due to errors in the population-based phasing. Since Nanopore reads are especially prone to errors in homopolymer regions, we investigated the effect of excluding all SNVs in such regions from phasing (see **Methods** for a precise definition). This resulted in a decrease in the number of established phase connections from 97.5% to 91.7% for Patient1 and from 96.5% to 91.1% for Patient2 (**Figure 5B**) and a decrease in the switch error rate with respect to the pedigree-based phasing from 1.4% to 0.9% in Patient1, see **Figure 5C**. This shows that switch errors are indeed often found at such homopolymer sites and that masking those sites significantly reduces switch error rates at the expense of only a moderate reduction of phased variants.

**Figure 5.**
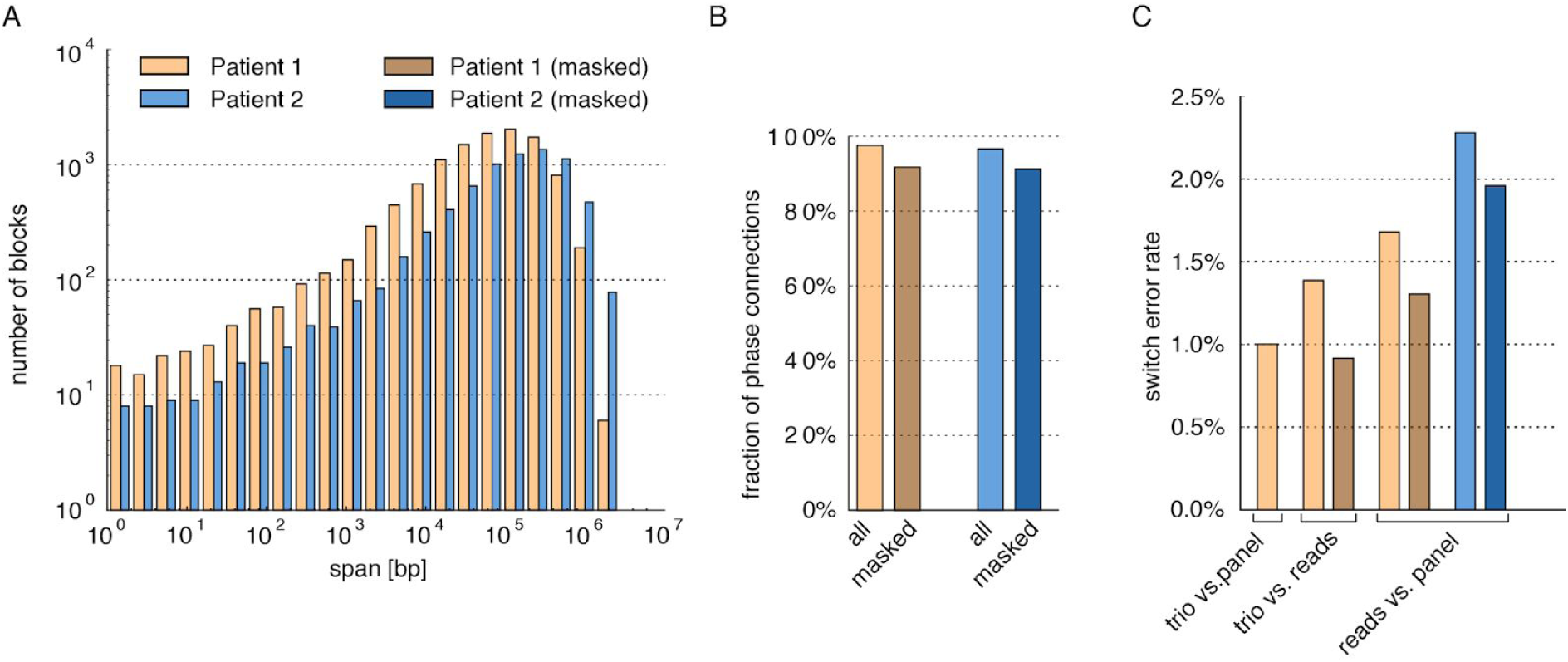
Performance of SNV phasing.

(A) Distribution of phased block lengths resulting from read-based phasing by WhatsHap. Patient1 and Patient2 are shown in brown and blue, respectively. (B) Fraction of phase connections (i.e. pairs of consecutive SNVs phased with respect to each other) established in the two patients and with/without masking repeats (light/dark colors). (C) For Patient1, switch error rates of all pairs of trio-based (PBT), population-based (ShapeIt), and read-based (WhatsHap) phasing are shown. For Patient2, where no family data is available, read-based phasing is compared to population-based phasing.

### Efficient phasing of genome-wide structural variations using nanopore reads

While structural variation has recently been integrated in larger population genetic reference panels, which enables their imputation for genetic association studies ^18,33^, building these panels often requires statistical phasing approaches, which drop accuracy for low allele frequency SV sites. Read-based phasing of SVs using long reads will enhance our ability to include SVs in high-quality reference panels, where structural variation is still underrepresented ^18^.

We apply the same methodology as above (i.e. used for phasing chromothriptic breakpoints) to evaluate genome-wide SV phasing accuracy. A total of 3.8M Nanopore reads overlapped one or more of the 1.7M genome-wide phase-informative SNPs. As estimated from reads overlapping at least 20 phase-informative SNPs, an average of 85.2% of the SNPs spanned by a read consistently support a particular phase assignment, which is in line with the reported error rate of MinION sequencing data (**Supplementary Figure 18**). A distinction between reads originating from paternal or maternal haplotypes can be readily made, particularly if reads overlap with multiple phase-informative SNPs (**Supplementary Figure 19**). We then selected a set of 2,389 heterozygous SVs that overlap between Manta (Illumina) and NanoSV (Nanopore) call sets. Each SV was assigned a phase and a phasing quality (**Methods**), by combining information from all phase-informative SNPs falling within the breakpoint-junction supporting reads and reference supporting reads respectively. In this way, we phased 1909 (78.7%) SVs and could assign 971 and 938 to paternal and maternal chromosomes, respectively. For the remainder of 480 SVs, spanning reads did not overlap any phase-informative SNP and therefore a phase could not be assigned to these SVs. Using the SV phasing produced by PBT as ground truth, our long-read based phasing of SVs had an accuracy of 98.5%.

## Discussion

In this work we demonstrate the feasibility of long-read sequencing of diploid human genomes on the MinION real-time portable nanopore sequencer. Given the long-read nature of the nanopore sequencing platform, we focused the analysis in this work on the detection of clinically relevant structural variations, a diverse category of genetic variation that is often causal to human genetic disease ^40^. A compelling motive for implementing long-read sequencing in the clinic is to diagnose patients with congenital phenotypes, such as intellectual disability ^41^. Hundreds to thousands of such patients are routinely screened annually for pathogenic SVs in clinical genetic centers, most often by copy number profiling or karyotyping. Although these methods are robust and relatively cost-efficient, they are not capable of mapping small or copy-balanced SVs, nor do they provide base-pair resolution accuracy, or the possibility to resolve complex SVs ^42^.

Here we show that MinION sequencing provides an attractive alternative approach for genome-wide structural variation detection, which could be implemented as a clinical screening tool. We were able to extract all known *de novo* breakpoint junctions for Patient1 (**Figure 1**), even at relatively low coverage. The long reads identified additional complexity for several breakpoint-junctions of Patient2. Moreover, 32% (29 vs 22) more chromothripsis breakpoint-junctions were detected with nanopore compared to short-insert Illumina sequencing. Our work also confirms previous data that revealed a substantial amount of novel SVs and indels discovered from PacBio long read sequencing of haploid human cells^29^. We observed that 33.3% of the high confidence set of SVs observed in the Nanopore data could not be found in matching Illumina sequencing data, despite the use of six different variant calling methods. Long sequencing reads thus enable a much more straightforward and homogeneous analysis of structural variation genome wide, while retaining a very high accuracy in the final set of variants.

Phasing of genotyped SVs - relevant for mapping disease associations - is commonly done using statistical methods or by employing family-relationships among sequenced individuals^18^. We here devised a computational strategy that allowed accurate phasing of SVs directly from the long nanopore reads using flanking heterozygous SNPs. Read-based phasing of SVs is advantageous particularly for classes of SVs with a low population frequency and for *de novo* variations. This is exemplified by the evidence provided here for the paternal origin of all *de novo* breakpoint-junctions in Patient1, whereas previous work on chromothripsis has not provided robust suport for the parental origin of chromothripsis.

If nanopore data quality improves at a similar pace as we observed during recent past (**Supplementary Figure 4**), SNV calling and genotyping may be directly performed based on the nanopore reads. Even though our data are of relatively low coverage, we were already able to obtain a good genotype concordance (96%) with the Illumina based pipeline, for existing SNV calls in Patient1 (data not shown). SNV calling combined with accurate phasing, as we demonstrated here, will enable nanopore-only genetic variation discovery and phasing.

The future of human genome sequencing will involve a shift towards longer sequencing reads that facilitate personal genome assemblies and alter the way we deal with genetic variation discovery and representation ^43^. Efforts to obtain full-length haplotype resolved chromosomal sequences are continuously advancing and the combination of multiple long-range sequencing and mapping approaches have recently led to diploid human genome assemblies with contig N50 size of well over 10Mb^13,44^. We have not attempted a full human genome assembly using the nanopore reads in this work. However, we were able to separate reads by haplotype, which formed the basis for a reconstruction of the long-range structure of chromothripsis rearrangements. Such information is essential for interpretation of clinical phenotypes ^45^.

A drawback of current short-read genome sequencing technology is the need for high capital investment, which often leads to sequencing infrastructure being located in dedicated sequencing centers. This is associated with a complex logistic workflow and relatively long turnaround times. Our results show that such limitations can be overcome by the use of portable nanopore sequencing technology. Since the start of this project in April 2016, we have seen a tenfold increase in throughput per MinION sequencing run (**Supplementary Figure 1**) and an increase in sequencing quality to 90% accuracy for high output 1D runs (450b/s). In practice this means that 10x coverage of the human genome can be reached using 10-15 MinION flowcells at a cost of 5,000$ to 8,000$ within one week of overall sequencing time.

This work provides a glimpse into the potential of long-read, real-time and portable sequencing technology for human genomics research and clinical application. Creating larger catalogues of SVs, in complex repeat regions and segmental duplications, is a particular challenge in the coming years. We foresee that population-scale genome sequencing by nanopore or other long read technology will facilitate such discoveries, leading to further understanding of the role of SVs in the human genome in general and genetic disease in particular.

## Methods

### Sample source

The DNA for human genome sequencing in this study was obtained from two patients with congenital abnormalities and the parents. Informed consent for genome sequencing and publication of the results was obtained from all subjects or their legal representatives. The study was approved by Institutional Review Boards of San Luigi University Hospital and Brigham and Women's Hospital and Massachusetts General Hospital. Both patients have been previously described by Redin et al ^16^.

### DNA extraction

DNA of Patient1 was obtained from either peripheral blood mononuclear cells (PBMCs) derived from blood and from renal epithelial cells obtained from urine. Renal cells were cultured up to 8 passages as reported previously ^46^. Cells were harvested after reaching confluency by trypsinization with TrypLE Select (Thermo Fisher Scientific) and centrifugation at 250g for 5 minutes. DNA from the parents was obtained from PBMCs. PBMCs were collected by a ficoll gradient. In brief, the blood was diluted 4x with phosphate buffered saline (PBS). Subsequently 13 mL of Histopaque(®)-1077 (family 1; Sigma-Aldrich 10771-500ML) was added per 35 mL of diluted blood. The resulting mixture was centrifuged at room temperature for 20 minutes at 900 x g, followed by recovery of the PBMC layer. PBMCs were washed twice using PBS, centrifuged at 750 x g for 5 minutes and resuspended in PBS with 50% DMSO. For Patient2, DNA was obtained from a lymphoblastoid cell line, which has not been tested for mycoplasma contamination. The cell line was authenticated by whole genome sequencing. DNA extraction from cultured cells and PBMCs was performed using DNAeasy (Qiagen) or Genomic-tip (Qiagen) according to manufacturer’s specifications with exclusion of vortexing to maintain DNA integrity.

### Nanopore library preparation and sequencing

Isolated DNA was sheared to ∼10-20kb fragments using G-tubes (Covaris). Subsequently, genomic libraries were prepared using the Oxford Nanopore Sequencing kit (SQK-MAP006 for R7 or SQK-NSK007 for R9), the Rapid library prep kit (SQK-RAD001) or the 1D ligation library prep kit SQK-LSK108. A 0.4x (instead of 1x) ampure cleanup was introduced after the FFPE DNA repair and the end-repair steps in the protocol to ensure removal of small DNA fragments. Genomic libraries were sequenced on R7.3, R9 and R9.4 flowcells followed by base-calling using either Metrichor workflows or MinKnow software. For Patient2 we introduced a DNA size selection step prior to library preparation using the Pippin HT system (Sage Science).

### Illumina whole genome sequencing

Genomic DNA of the patients and the parents was sheared to 400-500bp fragments using the Covaris. Subsequently, genomic libraries were prepared using the nano library preparation kit. Genomic libraries were sequenced on an Illumina HiSeq X instrument to a mean coverage depth of ∼30x.

### Nanopore data mapping

FASTQ files were extracted from base-called nanopore sequencing data using Poretools version 0.6.0 ^47^. Subsequently, fastq files were used as input for mapping by LAST (version 744)^17^, against the GRCh37 human reference genome. Prior to mapping the full dataset, we used the *last-train* function to optimize alignment scoring parameters using a sample of 1200 nanopore reads. Nanopore sequencing data were also mapped using BWA-MEM with the -x ont2d option. Nanopore 2D runs can produce 2D sequence reads, i.e. data where both forward and reverse reads of a DNA duplex are collapsed into a single sequence read, which produce three sequences in a fastq file, termed 1D template, 1D complement and 2D. Therefore, we filtered the LAST and BWA BAM files by only retaining one of these three “versions” for each read based on the following order of preference: 2D > 1D template > 1D complement.

### Illumina data mapping

Illumina HiSeq X ten data were mapped to the reference genome using BWA-0.7.5a using BWE-MEM -t 12 -c 100 -M -R. Reads were re-aligned using GATK IndelRealigner ^48^ and deduplication was performed using Sambamba markdup ^49^. Short indels and SNPs were genotyped using GATK HaplotypeCaller, jointly for the Patient1 trio and individually for Patient2.

### NanoSV algorithm

The NanoSV algorithm developed here (https://github.com/mroosmalen/nanosv) uses LAST BAM files as input. We did not use BWA-MEM alignments as NanoSV input, because the reads are not always split in non-overlapping segments. More precisely, we observed that the following two (oversimplified) CIGAR strings may be produced, for two aligned segments originating from the same sequencing read: 6M4S and 4S6M respectively. This observation was not further investigated for the purpose of this project.

NanoSV uses clustering of split reads to identify SV breakpoint-junctions. In a first step, all mapped segments of each split read are ordered based on their positions within the originally sequenced read. The aligned read may contain gaps between its aligned segments, i.e. parts of the read that do not align anywhere on the reference genome, for example due to insertions (**Supplementary Figure 8**, **Supplementary Figure 9**) or simply due to low quality sequencing.

Let tuple x = (c,s,e,k) describe an aligned sequence segment, where the chromosome and genomic start and end coordinates of the segment are specified by c, s and e respectively, and the mapping orientation by k ∈ {+,-}. The coordinates s and e represent that start (lowest) and end (highest) coordinate of the mapped segment on the reference genome. Now, read R_i_ can be described in terms of the ordered list of aligned segments and alignment gaps X_i_ = [u_1_, x_1_, u_2_, x_2_, u_3_, …, x_N_, u_N+1_], where the ordering is determined based on their occurrence in the read, u is the gap (i.e.: unaligned sequence preceding segment x) and N is the total number of aligned segments for read R. Alignment gaps are defined as read segments that are either unaligned or segments that fail to reach the mapping quality threshold Q_1_ (default: 20). The size of an unaligned segment is denoted as |u|, and can be zero in case two adjacent segments align successfully.

Any two consecutive aligned segments [x_n_, u_n_, x_n+1_] in a read define a candidate breakpoint-junction. We further aggregate evidence from different reads supporting the same candidate breakpoint-junction. This is achieved by clustering all candidate breakpoint-junctions that have the same orientation and have start and end coordinates that are in close genomic proximity. In order to facilitate clustering of reads that cover the same breakpoint-junction but that map to opposite strands of the reference human genome, order and orientation of the aligned segments is reverse complemented if for the genomic coordinates {p,q} mapping to the two closest bases of segments x_n_ and x_n+1_, respectively, within a given sequence read R_x_, at least one of the following conditions is met:

1. p and q are on the same chromosome and p-q>0
2. p and q are on different chromosomes and p has a higher chromosome number

The clustering is initialized by assigning each pair of consecutive aligned segments [x_n_, u_n_, x_n+1_] to a separate cluster. The resulting clusters are then recursively merged. Any two clusters (C_x_ and C_y_) are merged if and only if, there exists a candidate breakpoint-junction tuple (x_n_, x_n+1_) ∈ cluster C_x_ and a candidate breakpoint-junction tuple (y_m_, y_m+1_) ∈ cluster C_y_, such that the following conditions are met:

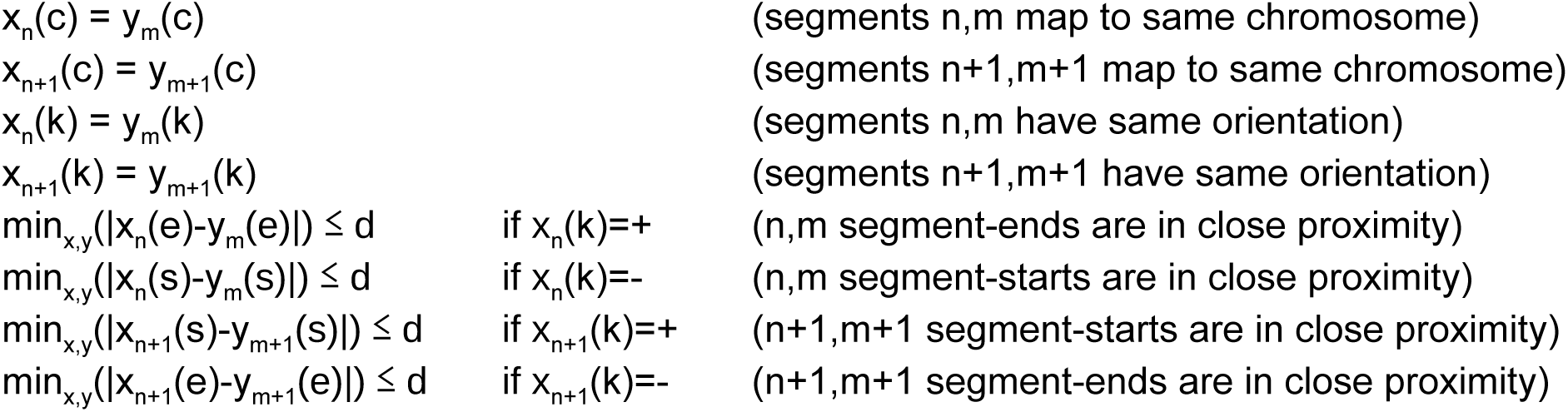

Where d is the threshold that we set for the maximum distance between the alignment coordinates of two segments if they are to be considered as supporting the same breakpoint-junction (default: 10 base-pairs). Recursive clustering continues until no more clusters can be merged. Each final cluster represents one candidate SV, which is described by tuple b = (c_1_,c_2_,p_1_,p_2_,k_1_,k_2_,g), with p_1_, p_2_ the medians of the start and end coordinates of all candidate breakpoint junctions contained in the cluster, c_1_, c_2_ the chromosomes associated to these coordinates and k_1_, k_2_ the orientation of the breakpoint-junction. Finally, the gap size g denotes the median size of the unaligned segments u_n_, between the two consecutive aligned segments x_n_ and x_n+1_ of all the tuples within the respective cluster.

A true SV is called when a candidate SV is supported by more than T reads (default: 2). Moreover, SVs with median mapping quality of the supporting reads not exceeding Q_2_ are still reported, but flagged as “MapQual” in the VCF FILTER field. SV-types can be determined from tuple b. Breakpoint-junctions where c_1_ and c_2_ point to different chromosomes are considered interchromosomal SVs (e.g. chromosomal translocations), which can have one of four possible orientations (3’to3’, 3’to5’, 5’to5’, 5’to3’). Similarly, breakpoint-junctions where c_1_ and c_2_ point to the same chromosome are intrachromosomal SVs, which can have one of four possible orientations (inversion type=3’to3’ or 5’to5’, deletion/insertion type=3’to5’, tandem duplication type=5’to3’). Insertions and deletions are discerned based on the relation between the gap size, g, and the reference-length l=|p_1_-p_2_|, where an insertion is called if g>l and a deletion is called when g=<l (**Supplementary Figure 9**).

We only consider two possible alleles for each SV candidate (present = ALT/absent = REF). The reads supporting the alternative allele contain the segments constituting the breakpoint-junction cluster. We consider as supportive of the reference allele all reads for which there is an aligned segment crossing one of the ends of the breakpoint junction (or both). More formally, a read is defined as crossing a breakpoint if it contains at least one aligned segment x_n_ for which holds: (p_1_ - x_n_(s) > 100 ∧ x_n_(e)-p_1_ > 100) ∨ (p_2_ - x_n_(s) > 100 ∧ x_n_(e) - p_2_ > 100). Reads not supporting the reference allele according to this definition are ignored. SV genotypes (homozygous alternative, heterozygous, homozygous reference, not-called) are assigned based on a Bayesian likelihood similar to the one used (and formally defined) by the SVTyper ^23^. SV calls are reported in VCF format following the VCF standards as maintained by samtools specifications ^50^. To facilitate reporting of complex SV types, such as inversions or reciprocal translocations, individual breakpoint-junctions that bridge the same chromosomal regions, but are opposite in orientation (e.g. 3’to3’ and 5’to5’ for inversions), are linked using an identifier.

### Nanopore data SV calling

We run NanoSV on the Nanopore data of each patient using the default parameters: “-t 8 -s 10 -p 0.70 -m 20 -d 10 -c 2 -f 100 -u 20 -r 300 -w 1000 -n 2 -q 80 -i 0.80 -g 100 -y 20”. We discarded all sites where the alternative allele count was 0 in the resulting genotype (i.e.: HOM_REF) and further filter the resulting call sets for SVs tagged as “Cluster”. The “Cluster” VCF INFO-field tag is added to all SV calls which lie inside a 500 base-pair region containing three SVs or more. These clusters of SVs are most likely either sequencing errors or located in highly repetitive and/or decoy regions of the human reference. We used Lumpy^23^ and Sniffles^24^ (specifically designed for Oxford Nanopore and Pacific Biosciences data) to call SVs in both samples using BWA-MEM alignments (instead of LAST alignment, as per requirement of the respective callers) of the same data and settings that match our own (liberal) NanoSV settings as closely as possible, as follows. For Lumpy: “-mw 2 -tt 0 -e”, requiring that at least one read supports each candidate breakpoint and clustering breakpoints within 10 base-pairs (back_distance=10). For Sniffles: “-s 1 --max_num_splits 10 -c 0 -d 10” ^24^. At the time of our analysis SVTyper was not supporting Nanopore reads (i.e. it required paired end reads), therefore we considered the Lumpy, ungenotyped, SV candidate sites as final calls for all subsequent analyses/comparisons.

### Random Forest variant filtering model

We trained a random forest (RF) model that we subsequently used to filter out false positive SV calls from our Nanopore data, such that we obtained a high precision set of variants for downstream analysis. The training data for our model consists of 191 true positive (TPs) SVs and 90 false positives (FPs). These 281 training data SVs are the highest confidence variants (TPs or FPs) resulting from the manual curation of 700 variants from the initial dataset, selected to span the whole SV size range and all SV classes. The manual curation was performed by two experts independently and only the sites where both experts made the same call were used. Out of the 191 TP SVs, 107 variants were also tested in one of the 3 initial validation rounds (see below) and in 89% of cases the validation call matches our manual curation; Similarly, 42 out of the 90 FP variants were also tested through validation and 69% of these labels are concordant.

The features supplied to the RF model are (where side1 and side2 refer to the lowest and highest genomic coordinates of a breakpoint-junction, respectively):

‐ Mapq1: average mapping quality over all reads supporting side1 of the breakpoint junction
‐ Mapq2: average mapping quality over all reads supporting side2 of the breakpoint junction
‐ Pid1: average percent identity (i.e.: to the reference) over all reads supporting side1 of the breakpoint junction
‐ Pid2: average percent identity (i.e.: to the reference) over all reads supporting side2 of the breakpoint junction
‐ Cipos1: genomic distance from the median start position of the SV to the lower bound of its associated confidence interval
‐ Cipos2: genomic distance from the median start position of the SV to the upper bound of its associated confidence interval (i.e.: confidence interval width = cipos1 + cipos2)
‐ Plength1: average proportion of the aligned segment (i.e.: relative to the entire read length), across all segments supporting side1 of the breakpoint junction
‐ Plength2: average proportion of the aligned segment (i.e.: relative to the entire read length), across all segments supporting side2 of the breakpoint junction
‐ Ciend1: genomic distance from the (median) end position of the SV to the lower bound of its associated confidence interval
‐ Ciend2: genomic distance from the (median) end position of the SV to the upper bound of its associated confidence interval (i.e.: confidence interval width = ciend1 + ciend2)
‐ totalCovNorm: depth coverage summed across both ends of the breakpoint junction, divided by the average depth of coverage across the sample
‐ Vaf: percentage of the reads spanning either end of the breakpoint junction that support the variant allele (i.e.: the presence of a breakpoint junction)

The precision-recall curve of the model, and its 95% confidence interval, displayed in **Supplementary Figure 16** is derived from 100 bootstrapping runs where the whole training data was split into 80%-20% train-test subsets. The optimal operating point was chosen at 92% precision and 83% recall.

The model trained on the whole training data was then applied to the whole set of 8,667 SVs to produce a high confidence set of 4,587 SV calls, for Patient1.

### Illumina data SV calling

SV calling for Illumina data was done using Manta^20^, Delly^19^, FREEC^31^, Mobster^30^ and Pindel^32^. For Manta we used version 0.29.5 with standard settings, for Delly we used version 0.7.2 with “-q 1 -s 9 -m 13 -u 5”, for FREEC we used version 7.2 with window=1000, for Mobster we used version 0.1.6 with standard settings (Mobster properties template), for Pindel we used version v0.2.5b8 with standard settings and excluding regions represented by the UCSC GRCh37 gap table (https://genome.ucsc.edu) using the -c option. Homozygous reference calls (genotype = 0/0) were omitted from the call sets for each of these tools.

### PCR, primer design and SV validations

Primers for breakpoint-junction validation were designed using Primer3 software^51^. Breakpoint-junction coordinates and orientations were used as input for primer design. Amplicon sizes varied between 500-1000bp. PCR reactions were performed using AmpliTaq gold (Thermo Scientific) under standard cycling conditions. PCR products were sequenced using MiSeq (TruSeq library preparation, Illumina), Sanger sequencing (Macrogen) or MinION Nanopore sequencing (2D library preparation and sequencing).

We perform extensive and heterogeneous validation on the SV calls of Patient1, in order to obtain a thorough and reliable characterization of our dataset and an informative comparison to standard approaches. We first randomly selected 384 NanoSV candidate calls (uniformly distributed across the observed size-range of SVs) from the call set of Patient1 and performed validation with Illumina MiSeq. We further selected 400 candidate calls (uniformly distributed across the observed size-range of SVs) exclusively from the Nanopore specific SV calls and validated them. Deep coverage Nanopore sequencing was used for this second round of validation, under the assumption that a long-read accessible only set of variants would be less likely to validate using the short read Illumina sequencing. A third round of validation was performed, also by Nanopore deep coverage sequencing, on a set of 192 non-random variants; namely, 96 variants were expected to be true positive SV calls and 96 false positive SV calls, as of an initial attempt to build a discriminative model. Upon inspection of these validation results, SVs falling within homopolymer stretches (see above) and/or short tandem repeats (UCSC tandem repeat table) were considered unreliably genotyped (i.e. even in the validation data) and were subsequently discarded from the dataset altogether (see main text - **Results**).

All of the above three rounds of validation are thus restricted to the sites that fall outside homopolymers and/or short tandem repeats and SVs for which we did not obtain a specific PCR product are discarded. This is the subset that is referred to as validation data throughout the text, when evaluating precision and it consists of 273 SVs (193 true positives and 80 false positives). The dataset used to train the Random Forest for the post-calling filtering step (191 true positives and 90 false positives) is a subset of the validation data assays, augmented with 83 more SVs, randomly selected from the initial calls and manually evaluated for validation status. The 75 SVs used as test data for the Random Forest model are all a subset of the deep sequencing validation assays.

A structural variant was considered validated as a true positive if there exists an SV call, in the validation SV call set, that overlaps (in the meaning described below) the original SV validation candidate. The validation SV call set is produced similarly to the initial analysis, where Manta is used for genotyping SVs in the MiSeq validation data and NanoSV is used for the Nanopore data respectively, with the note that deep coverage (i.e.: ∼1,000 for MiSeq and Nanopore runs) enables accurate genotyping.

### Calculating overlap between SV datasets

To calculate the intersection between SV call sets, we considered each SV call as a set of breakpoint-junction start and end coordinates s and e, and orientation k. For any SV call i, each breakpoint-junction coordinate (s_i_ and e_i_) is the median of an associated confidence interval, (s_i,l_,s_i,h_) and (e_i,l_,e_i,h_) respectively, as derived from the evidence cluster C_i_. SV calls i and j are overlapping if the confidence intervals of their start and end coordinates are closer together than 101 bp. For SVs smaller than 1000bp (excluding insertions), we additionally required that SVs overlap each other with a reciprocal overlap of at least 70%, otherwise, considering the 100 base-pair margin that we use when comparing breakpoint junction borders, different SVs that happen to be in genomic close proximity may, incorrectly, be considered the same event.

### GC bias

The GC content (i.e. percentage of guanine or cytosine bases within a certain DNA sequence) was computed for 100,000 5kb intervals of the reference genome (build GRCH37). These intervals were chosen such that they do not overlap sequencing gaps in the reference, as defined in the UCSC table browser (https://genome.ucsc.edu/cgi-bin/hgTables), including telomers, centromeres and other gaps. The average depth of coverage across each interval was then computed from the HiSeq alignment data and the Nanopore alignment data respectively (stratified by sequence reads tagged as “passed” and “failed” by the Metrichor basecalling for Patient1). The GC content was binned into 30 uniformly spread bins, between the minimum and the maximum GC content derived from the data. Six GC-content bins were discarded - i.e. those where GC-content < 0.26 or GC-content > 0.66 - as too few sampled intervals fall within these bins and a coverage distribution cannot be robustly estimated (i.e.: 1 - 18 intervals per bin, **Supplementary Figure 6**).

A linear regression model with average coverage as the dependent variable and GC-content as the independent variable was trained, in order to quantify the GC bias of the two sequencing technologies, respectively. The average coverage values were normalized (0 mean, 1 variance) for Illumina and Nanopore data respectively, because of the different sequencing average depth of coverage, such that the regression coefficients for the two technologies be comparable (i.e. the resulting regression coefficients express the number of standard deviations that the coverage varies, per GC content percentage).

### Genetic phasing of SNPs, indels and SVs from Illumina sequencing data

We used the Illumina whole genome sequencing data of Patient1 and both its parents to obtain a high confidence set of phased genotypes (including SNPs, short indels and SVs), against which we subsequently evaluated the Nanopore data analysis. We used GATK PhaseByTransmission (PBT) ^26^ to correct genotypes based on trio information and to obtain deterministic phasing for most loci. PBT settings were: “-prior 0.000001 -useAF GT -af_cap 0.0001”. The PBT-phased SNPs were used to evaluate the genome-wide read-backed phasing from Nanopore data as well as for phasing the Nanopore reads and the PBT-phased SVs were used to evaluate the Nanopore read-backed phasing of the SVs (i.e.: evaluation was limited to the SVs detected in both Nanopore and Illumina data). PBT was run with a de novo mutation prior of 10e-6 and supplied with the population allele frequencies of 1000 genomes Phase 3 European population.

### Nanopore read-based phasing of SNVs using Whatshap

For both patients, all bi-allelic heterozygous SNVs were phased from the aligned Nanopore reads using WhatsHap (version 0.13+21.g45bd7f8, ^37,38^) with realignment mode enabled.

That is, reads were realigned against reference and alternative alleles at variant sites, which is critical for phasing performance of noisy long reads ^38^. For comparison purposes, we used SNV genotypes to obtain a population-based phasing with respect to the 1000 Genomes phase 3 ^39^ reference panel by running ShapeIt with default parameters. We excluded from the comparison all variants that fell within homopolymer runs longer or equal to 5 base-pairs, due to both genotyping accuracy, but mostly because of Nanopore’s known drop in sequencing accuracy for longer homopolymer sequences. The homopolymer bed-track used was computed genome-wide, incorporating a one base-pair border around the homopolymer, such that relatively frequent sequences of the form “XXXXXYZZZZZ” be merged into one homopolymer region for the final result.

### Phasing of nanopore reads and SVs

Individual nanopore reads from Patient1 were phased using a set of 1.7M heterozygous SNPs that were genetically phased by GATK PBT^26^. Individual nanopore reads were phased using the genetically phased SNPs by determining the basecall and corresponding basequality at each SNP position within each read. Let b(i) and q(i) be the basecall and associated quality value for some SNP *i* in some read under evaluation. Further let BM(i) and BP(i) be the maternal and paternal alleles respectively (i.e.: as phased by PBT), for SNP *i*. The information from all SNPs spanned by a read is then aggregated and the likelihood that read r originates from the paternal or the maternal haplotype respectively is computed:

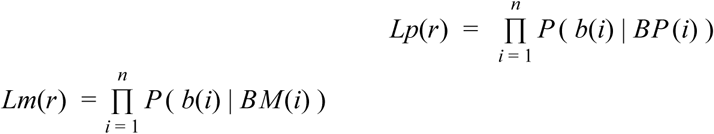

Where *n* is the total number of SNPs that read *r* overlaps and

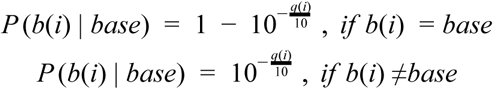

Is the probability that a read supports a specific phased allele at a SNP. The likelihoods that the SV resides on the paternal or the maternal haplotype respectively are then computed:

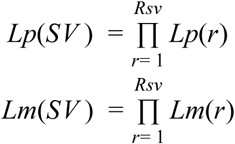

Where *Rsv* denotes the set of all reads supporting the breakpoint junction. The two likelihood scores are then transformed to probabilities (i.e.: normalized to sum up to 1) and phase for the set of breakpoint-junction supporting reads is assigned as indicated by the highest likelihood score. Phase is assigned identically to the set of reference-supporting reads spanning the breakpoint junction.

An SV is then considered phased if the two phases, for the set of breakpoint supporting reads and reference supporting reads respectively, correspond to different parental haplotypes and the (phred scaled) phasing posterior quality is defined as:

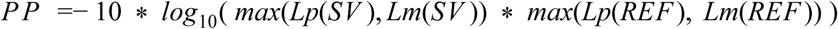

### Construction of chromothripsis structure using reference-based nanopore read overlap

To obtain evidence for the long-range structure of the chromothripsis breakpoint-junctions in Patient1, we first extracted the set of (aligned) Nanopore reads that span the chromothripsis regions on chromosomes 1, 7, 8 and 9. Separation of reads by phase was done as described above. Each mapped segment was ordered by left genomic mapping coordinate of each segment to produce an ordered list of segments L={s(1), s(2), …, s(n)}. Nodes were constructed from the ordered segments by requiring overlap between consecutive segment. Let i and j represent the start (left) and end (right) coordinate of segment s(1) and s(2). In order for s(1) and s(2) to be assigned to the same node, we required (s(1)_j_-s(2)_i_) >= 20bp. A segment s(n) is also assigned to the same node as the previous segment if the overlap with the previous segment in the ordered list L is <20bp, but only if s(n) overlaps 20bp with at least two earlier segments in L.

In a subsequent step individual nodes are connected based on overlapping read names. Consider the following reads (r), segments (s) and nodes (m):

- r(1)-s(1)-> m(1)
- r(2)-s(1)-> m(2)
- r(1)-s(2)-> m(2)
- r(2)-s(2)-> m(3)

The path through the nodes is considered as m(1)-m(2)-m(3). The number of individual read names that connect nodes is required to be at least 2.

Using the above algorithm, individual breakpoint junctions were connected together, providing support for the order of the joined segments in the chromothripsis chromosomes of Patient1.

### Assembly of Nanopore sequencing data

Nanopore reads of Patient1 were separated into three bins by phase, as described above. The reads that were assigned a paternal phase and the unphased reads were used as input for *de novo* assembly using Miniasm^28^, with settings: minimap -S -w 5 -L 100 -r 500 -m 0 and miniasm -c 1 -m 100 -h 20000 -s 100 -r 1,0 -F 1. The contigs outputted by Miniasm were aligned to the human reference genome (GRCh37) using LAST, with settings: -s 2 -T 0 -Q 0 -p [*last_parameters*]. The *last_parameters* were obtained as described above. LAST aligments (SAM format) were processed by custom scripts to evaluate the presence of chromothripsis segments from Patient1 based on chromosomal coordinate overlap.

**Competing interest statement**

WK and JdR have received financial compensation for travel and accommodation expenses to speak at an Oxford Nanopore-organised meeting.

## Acknowledgements

We thank the Bioinformatics Expertise Center of the UMC Utrecht for setting up part of the computational infrastructure and software to analyze nanopore sequencing data. This work was supported by funds from the Utrecht University to implement a single-molecule sequencing facility. MCS is supported by VIDI grant 91712354 from the Dutch organization for scientific research (NWO-ZONMW). We thank Eleonora Di Gregorio and Alfredo Brusco for their contribution to the identification of the complex chromosomal rearrangement in Patient1.

## Author Contributions

SM, GP, DG, GM, JK, MET provided access to patient cells and DNA. IR and WPK generated nanopore sequencing data. EB, JK and EC provided Illumina sequencing data. MCS, MJR, MN, JL, JEVI, TM, JR and WPK performed nanopore data analysis. MCS, WPK, TM and JR wrote the manuscript. All authors contributed to the final version of the manuscript.

